# Sensor NLR immune proteins activate oligomerization of their NRC helper

**DOI:** 10.1101/2022.04.25.489342

**Authors:** Mauricio P. Contreras, Hsuan Pai, Yasin Tumtas, Cian Duggan, Enoch Lok Him Yuen, Angel Vergara Cruces, Jiorgos Kourelis, Hee-Kyung Ahn, Chih-Hang Wu, Tolga O. Bozkurt, Lida Derevnina, Sophien Kamoun

## Abstract

Nucleotide-binding domain and leucine-rich repeat (NLR) immune receptors are important components of plant and metazoan innate immunity that can function as individual units or as pairs or networks. Upon activation, NLRs form multiprotein complexes termed resistosomes or inflammasomes. Whereas metazoan paired NLRs, such as NAIP/NLRC4, activate into hetero-complexes, the molecular mechanisms underpinning activation of plant paired NLRs, especially whether they associate in resistosome hetero-complexes is unknown. In asterid plant species, the NLR required for cell death (NRC) immune receptor network is composed of multiple resistance protein sensors and downstream helpers that confer immunity against diverse plant pathogens. Here, we show that pathogen effector-activation of the NLR proteins Rx (confers virus resistance) and Bs2 (confers bacterial resistance) leads to oligomerization of the helper NLR NRC2. Activated Rx does not oligomerize or enter into a stable complex with the NRC2 oligomer and remains cytoplasmic. In contrast, activated NRC2 oligomers accumulate in membrane-associated puncta. We propose an activation-and-release model for NLRs in the NRC immune receptor network. This points to a distinct activation model compared to mammalian paired NLRs.

## Introduction

A class of intracellular immune receptors, known as NLRs (nucleotide-binding domain and leucine-rich repeat receptors), are key components of the innate immune systems of plants and metazoan. NLRs play an important role in mediating pathogen recognition and subsequent immune responses (Duxbury *et al*, 2021; Jones *et al*, 2016). In plants, NLRs can activate host defense by recognizing pathogen secreted virulence proteins, termed effectors. This recognition leads to immune signaling, often culminating in a form of programmed cell death known as the hypersensitive response (Jones & Dangl, 2006; Kourelis & Van Der Hoorn, 2018; Ngou *et al*, 2022). Similarly, metazoan NLRs are capable of sensing pathogen effectors and other classes of pathogen derived molecules, ultimately leading to a form of programmed cell death known as pyroptosis (Duxbury *et al*, 2016; Maekawa *et al*, 2011). Some plant and metazoan NLRs can function as single units, with one NLR protein mediating both effector/elicitor perception and subsequent downstream signaling. These are referred to as functional singleton NLRs (Adachi *et al*, 2019a; Adachi *et al*, 2019b). However, NLRs can also function as genetically linked receptor pairs or in higher order configurations that can include genetically unlinked receptor networks (Wu *et al*, 2017; Wu *et al*, 2018). In these cases, the sensing and signaling functions are uncoupled in two distinct proteins. One NLR acts as the pathogen sensor, requiring a second NLR which acts as a helper (or executor) to mediate immune activation and disease resistance (Adachi *et al*., 2019b; Feehan *et al*, 2020; Wu *et al*., 2018). Although much progress has been made in recent years regarding the biochemical mechanisms of how singleton NLRs activate and signal (Förderer *et al*, 2022; Ma *et al*, 2020; Martin *et al*, 2020; Wang *et al*, 2019a), our understanding of how paired and networked NLRs operate remains limited.

Plant and metazoan NLRs belong to the signal transduction ATPases with numerous domains (STAND) superfamily. They usually exhibit a modular, tri-partite structure with an N-terminal signaling domain, a central nucleotide binding domain and a C-terminal domain with superstructure forming repeats (Duxbury *et al*., 2021; Kim *et al*, 2016; Kourelis *et al*, 2021b). The N-terminal domains of NLRs can broadly be used to classify these receptors into distinct groups which, in plants, tend to also cluster together in phylogenetic analyses. Plant NLR N-terminal domains can be either coiled-coil-type (CC) NLRs, G10-type CC (CC_G10_) NLRs, RPW8-type CC (CC_R_) NLRs or toll/interleukin-1 receptor-type (TIR) NLRs, whereas metazoan NLRs usually exhibit either N-terminal PYRIN or caspase recruitment domains (CARD) (Kim *et al*., 2016; Kourelis *et al*., 2021b; Lechtenberg *et al*, 2014; Lee *et al*, 2021). The central nucleotide binding domain is the defining feature of NLRs and is typically a nucleotide-binding adaptor shared by APAF-1, plant R proteins and CED-4 (NB-ARC) domain in plants, while metazoan NLRs can have either an NB-ARC or a NAIP, C2TA, HET-E and TP1 (NACHT) domain. As for their superstructure forming repeats, these can be either leucine-rich repeats (LRR) or tetratricopeptide repeats (TPR) (Duxbury *et al*., 2021; Kourelis *et al*., 2021b).

While the general principles of how NLRs sense their ligands are well-understood (Kourelis & Van Der Hoorn, 2018; Ngou *et al*., 2022), our knowledge of the molecular mechanisms that underpin NLR activation and signaling are more limited. In the case of mammalian NLRs, activation leads to oligomerization and formation of higher order wheel-like complexes, termed inflammasomes. Inflammasomes ultimately recruit caspases which act as the final executors of programmed cell-death (Kim *et al*., 2016; Lechtenberg *et al*., 2014). In contrast, the mechanisms of plant NLR activation were not well understood until the recent elucidation of the structure of inactive and activated ZAR1, a conserved singleton CC-NLR from Arabidopsis (Adachi *et al*, 2022; Wang *et al*., 2019a; Wang *et al*, 2019b). The activation of ZAR1 upon recognition of its cognate effectors both *in vitro* and *in vivo* leads to its oligomerization and formation of a higher-order pentameric homo-complex analogous to the inflammasome and coined as the resistosome (Hu *et al*, 2020; Wang *et al*., 2019a; Wang *et al*., 2019b). Oligomerization based activation mechanisms have also been observed *in vivo* for the plant singleton CC-NLR RPP7 and the CC_R_-NLR helper NRG1.1 (Jacob *et al*, 2021; Li *et al*, 2020). More recently, the structure of the activated wheat CC-NLR Sr35 pentameric resistosome suggests that this activation strategy is likely evolutionarily conserved across plant CC-NLRs (Förderer *et al*., 2022). The activated complexes of CC-NLRs and CC_R_-NLRs act as Ca^2+^-permeable membrane-associated pores upon complex formation, an activity that is required for the hypersensitive cell death (Bi *et al*, 2021; Duggan *et al*, 2021; Förderer *et al*., 2022; Jacob *et al*., 2021; Ngou *et al*, 2021). Despite these advances, the molecular mechanisms of paired and networked plant CC-NLR activation are poorly understood. In the case of Pia (RGA4 and RGA5), immune signaling is activated through release of negative regulation (Césari *et al*, 2014). In contrast, the Pik-1 and Pik-2 pair is activated via receptor cooperation by forming a tri-partite complex with the pathogen effector (De la Concepcion *et al*, 2021; Zdrzałek *et al*, 2020). However, whether sensor and helper NLRs engage in heteromeric resistosome complexes is unknown.

Networked NLR immune signaling architectures present many advantages to plant and metazoan immune systems, likely contributing to their robustness and enhancing their evolvability in the face of highly adaptable pathogens (Wu *et al*., 2018). In mammals, multiple different NAIP sensor NLRs can perceive distinct immune elicitors and switch to an active conformation, contributing to immunity. Following activation, NAIPs require the helper NLR NLRC4 to mediate downstream signaling (Kofoed & Vance, 2011; Vance, 2015; Zhao *et al*, 2011). NAIP2 is one of these sensors. Upon perception of its cognate effector, NAIP2 initiates sensor-helper signaling via the formation of a heterocomplex with NLRC4 (Qu *et al*, 2012). This NAIP2/NLRC4 heterocomplex acts as a nucleation point for multiple NLRC4 monomers that leads to the formation of a NAIP/NLRC4 inflammasome with multiple additional NLRC4 units (Hu *et al*, 2015; Zhang *et al*, 2015). Similar networked signaling architectures have also been described in plants. In asterid flowering plants, a major phylogenetic cluster of CC-NLRs known as the NRC superclade comprises an immune receptor network with multiple sensor NLRs and downstream helper NLRs (Wu *et al*., 2017). This network mediates immunity against a wide array of plant pathogens including oomycetes, nematodes, aphids, bacteria and viruses and includes many well characterized and agronomically important sensor NLRs (Derevnina *et al*, 2021; Wu *et al*., 2017). All sensors in the NRC network signal, often redundantly, through a downstream hub of helper NRCs to mediate cell death and disease resistance. NRC helpers contain a key signature in the **α**-1 helix of their N-termini known as the MADA motif, which is crucial for mediating cell death. This motif is conserved in around 20% of angiosperm CC-NLRs and is functionally conserved between ZAR1, Sr35 and the NRCs, which suggests that the ‘death switch mechanism’ characterized for the ZAR1 resistosome may apply to NRCs as well (Adachi *et al*., 2019a; Förderer *et al*., 2022; Kourelis *et al*, 2021a). Considering how widespread and vital this immune network is for several crop species, developing a better mechanistic understanding of how it functions is critical. However, how sensor and helper NLR pairs communicate and initiate immune responses is not understood.

In this study, we explore the mechanism of activation of a sensor/helper CC-NLR pair. We selected the NRC-dependent sensor CC-NLR Rx as a model experimental system. Rx is an agronomically important sensor NLR from potato (*Solanum tuberosum)* that confers resistance to *Potato virus X* (PVX), a single-stranded RNA filamentous plant virus, by recognizing its coat protein (CP) (Bendahmane *et al*, 1999; Bendahmane *et al*, 1995; Grinzato *et al*, 2020). Prior to activation, Rx is held in an inactive state by intramolecular autoinhibitory interactions between its LRR domain and its CC and NB-ARC domains (Moffett *et al*, 2002; Rairdan & Moffett, 2006). Upon CP-triggered activation, Rx undergoes intramolecular rearrangements that include the release of LRR autoinhibition and the exposure of its NB-ARC domain, leading to its activation (Moffett *et al*., 2002). We previously showed that in order to mediate hypersensitive cell death and disease resistance, Rx and other sensors in the NRC network genetically require their downstream NRC helpers, with different sensors exhibiting different NRC helper specificities (Derevnina *et al*., 2021; Wu *et al*., 2017). Rx and the wild pepper (*Capsicum chacoense*) NLR Bs2, for example, can signal interchangeably via NRC2, NRC3 or NRC4. In contrast, the *Solanum bulbocastanum* NLR Rpi-blb2 which confers resistance to *Phytophthora infestans* strains carrying AVRblb2, can only signal through NRC4 (**Figure S1A**). However, the mechanisms by which sensor NLRs signal through NRCs are still not understood. Similar to the mammalian NAIP/NLRC4 system, Rx could be forming distinct Rx/NRC higher order hetero-resistosomes with each of its three NRC helpers, reminiscent of the NAIP/NLRC4 inflammasomes (Adachi *et al*., 2019b). Alternatively, plants may feature distinct activation mechanisms for paired and networked NLRs than those previously shown in mammalian paired systems. How activation of sensor NLRs translates into helper activation, immune signaling and disease resistance remains a fundamental question in plant immunology.

To dissect the biochemical mechanisms that underpin Rx and NRC activation, we established a resistosome formation assay using Blue Native polyacrylamide gel electrophoresis (BN-PAGE) by taking advantage of NRC proteins mutated in the MADA motif that are activated without triggering plant cell death. We demonstrate Rx-mediates oligomerization of its NRC2 helper in *Nicotiana benthamiana* following virus perception. Our data suggest that the activated NRC2 complex is an NRC2 resistosome that does not include Rx. Confocal live cell imaging and membrane fractionation assays reveal a sub-cellular shift in localization for NRC2 upon resistosome formation, moving from the cell cytoplasm to the plasma membrane to form membrane-associated punctate structures. This points to an activation-and-release model for sensor-helper signaling in the NRC network, whereby Rx can trigger NRC2 oligomerization without stably forming part of the activated helper complex. Notably, this model is distinct from the hetero-complexes shown for mammalian NLR paired systems, such as NAIP/NLRC4, implying that plant and metazoan NLR pairs exhibit different activation strategies.

## Results

### Activation of Rx with *Potato virus X* coat protein leads to oligomerization of its helper NRC2

Prior biochemical *in vivo* studies of activated NLRs have been hampered by the cell death response initiated upon receptor activation. We hypothesized that we could leverage N-terminal MADA motif mutations which abolish cell death induction without compromising resistosome formation (Adachi *et al*., 2019a; Duggan *et al*., 2021; Förderer *et al*., 2022; Hu *et al*., 2020). We previously showed that NRC2 is a MADA-type NLR with a conserved motif in its N-terminus, which is required for the execution of cell death and immune signalling. Making L9E, L13E and L17E mutations in the MADA motif of NRC4 completely abolishes NRC4 mediated cell death without affecting protein stability (Adachi *et al*., 2019a; Kourelis *et al*., 2021a). To this end, we created MADA mutations in NRC2 analogous to those previously made in NRC4, to generate an NRC2^L9/13/17E^ MADA motif mutant (NRC2^EEE^) (**Figure S1B**). Like the previously characterized NRC4 mutants, NRC2^EEE^ is unable to functionally complement hypersensitive cell death upon co-expression of effector-activated NRC2-dependent sensors in *nrc2/3/4 N. benthamiana* CRISPR mutant lines (**Figure S1C**).

We subsequently leveraged Rx and NRC2^EEE^ to determine the oligomeric state of the sensor and helper upon effector-triggered activation *in vivo*. To achieve this, we transiently expressed these proteins by agroinfiltration in the absence or presence of PVX CP in leaves of *nrc2/3/4 N. benthamiana* CRISPR lines using agroinfiltration and subjected total protein extracts to BN-PAGE assays (**Figure 1A**). In its inactive state NRC2^EEE^ appears as a fast-migrating band of ∼200 kDa, regardless of the presence or absence of Rx (**Figure 1B**). Upon activating the system by co-expressing Rx and NRC2^EEE^ together with C-terminally tagged CP-GFP, but not upon co-expression of free GFP as a negative control, NRC2 shifts to a slower-migrating, high molecular weight complex with two bands in the 720 to 1048 kDa range (**Figure 1B**). While we consistently observe this two-band pattern, with a lower molecular weight band of ∼750 kDa and a higher molecular weight band of ∼900 kDa, the ∼900 kDa band is more abundant. This shift to a higher molecular weight complex upon activation of the system is reminiscent of the ZAR1 resistosome formation documented *in vivo* (Hu *et al*., 2020), suggesting that NRC2 may exhibit a similar mechanism of oligomerization and resistosome formation upon activation. The formation of this higher order NRC2 complex is dependent on the presence of Rx, as in the absence of the upstream sensor, co-expression with CP does not result in oligomerization of NRC2^EEE^ or any other apparent change in molecular weight as compared to in the absence of CP (**Figure 1B**). We conclude that Rx is able and required to mediate NRC2 oligomerization upon CP-triggered activation.

**Figure 1:**
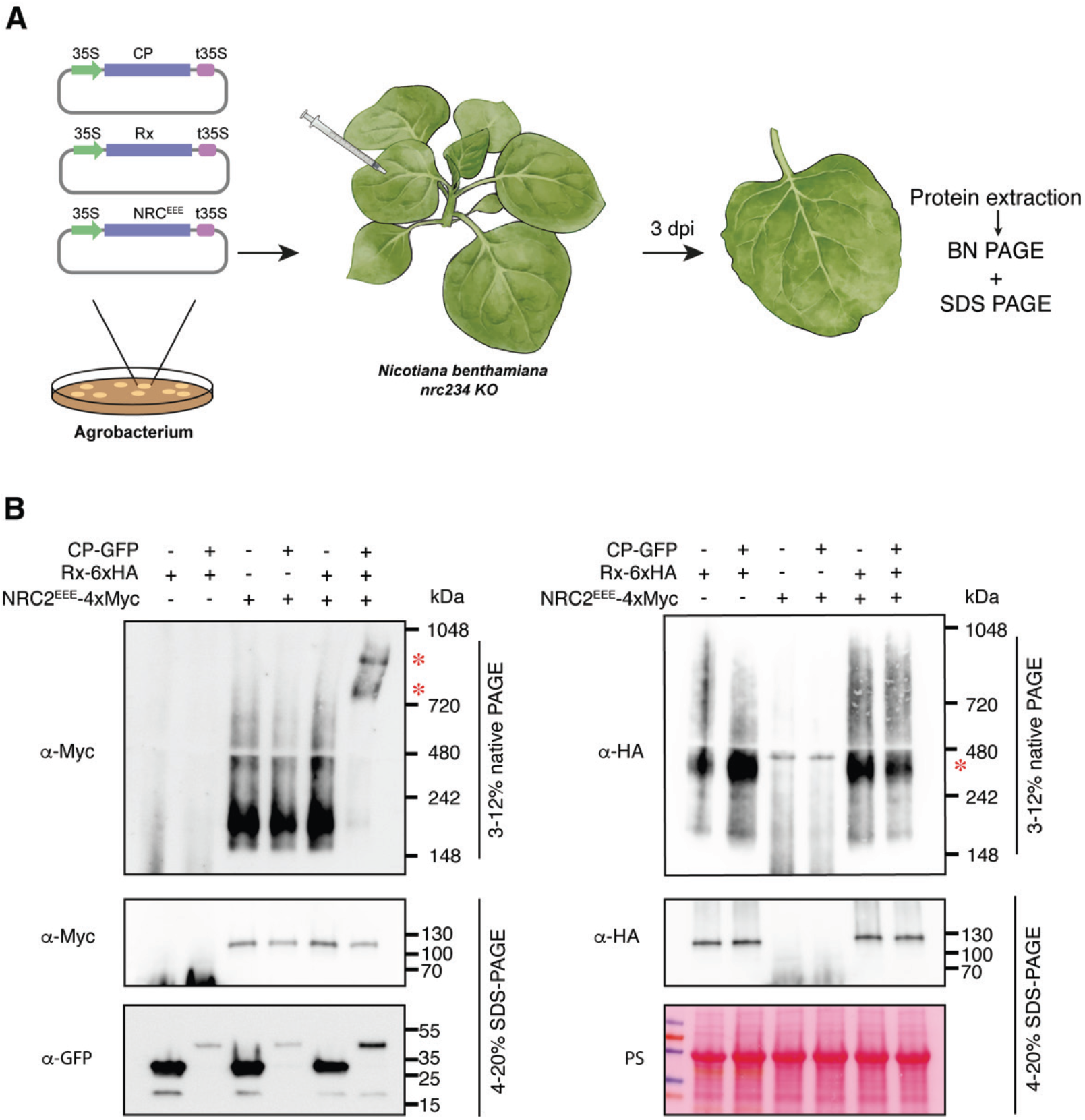
Activation of Rx with *Potato virus X* coat protein leads to NRC2 oligomerization. (**A**) Schematic representation of the experimental pipeline used. *Agrobacterium tumefaciens* strains were used to transiently express proteins of interest in leaves of *nrc/2/3/4 Nicotiana benthamiana* CRISPR mutant lines by agroinfiltration. Leaf tissue was harvested 3 days post-infiltration and total protein extracts were used for BN and SDS-PAGE assays. (**B**) BN-PAGE and SDS-PAGE assays with inactive and activated Rx-NRC2. C-terminally 6xHA tagged Rx and C-terminally 4xMyc-tagged NRC2^EEE^ were co-expressed with either free GFP or C-terminally GFP-tagged CP. Free mCherry-4xMyc and mCherry-6xHA fusions were used as controls in the treatments without NRC2 and Rx, respectively. Total protein extracts were run on native and denaturing PAGE assays in parallel and immunoblotted with the appropriate antisera labelled on the left. Approximate molecular weights (kDa) of the proteins are shown on the right. Red asterisks indicate bands corresponding to the activated NRC2 complex. Rubisco loading control was carried out using Ponceau stain (PS). The experiment was repeated three times.

### Rx does not oligomerize following activation by CP

In the BN-PAGE assays performed above, we observed that the sensor Rx constitutively migrates as a band of ∼400 kDa, regardless of its activation state (**Figure 1B**). Interestingly, the presence of this ∼400 kDa band does not depend on the presence of NRC2^EEE^ in *N. benthamiana nrc2/3/4* CRISPR mutant lines, suggesting that it is unlikely to be a preformed complex between Rx or any endogenous NRC2, NRC3 or NRC4. Considering that the size of a single C-terminally Rx-6xHA monomer is expected to be ∼115 kDa, we hypothesized that this ∼400 kDa band could correspond to a preformed Rx complex with additional host proteins. Rx could also potentially constitutively self-associate, forming a larger complex of multiple sensor units. We also did not observe any changes in size for Rx upon activation of the system with CP (**Figure 1B**). We were unable to detect any signal for Rx at a size matching the activated higher molecular weight NRC2^EEE^ complex described above, suggesting that Rx does not form a stable part of this activated NRC2 complex (**Figure 1B**). This implies an activation mechanism that differs from the hetero-oligomeric inflammasome previously reported for mammalian paired NLRs.

### Rx does not form a stable complex with NRC2

To further test the hypothesis that Rx does not form part of the activated NRC2 complex, we decided to use different molecular weight tags to study Rx-NRC2 interactions with size shifts. In previous experiments, we employed C-terminally tagged Rx-6xHA and NRC2^EEE^-4xMyc (from here onwards termed “light” versions) (**Figure 2A**). We then generated C-terminal tandem tag constructs to generate Rx-mCherry-6xHA and NRC2^EEE^-mCherry-4xMyc (termed “heavy” versions) (**Figure 2A**). We used these higher molecular weight versions to determine whether the addition of a larger molecular weight tag in one of the Rx-NRC2 system components could induce a shift in size of the other in BN-PAGE assays. We first confirmed that these new “heavy” versions retained the ability to correctly mediate hypersensitive cell death (**Figure S2**). When complementing leaves of *nrc2/3/4 N. benthamiana* CRISPR mutant lines with light NRC2^EEE^-4xMyc together with the heavy Rx-mCherry-6xHA, BN-PAGE assays showed no shift in size for NRC2^EEE^, either in the inactive or activated states, relative to light NRC2^EEE^-4xMyc co-expressed with light Rx-6xHA (**Figure 2B)**. Heavy Rx-mCherry-6xHA, in contrast, exhibited a shift in size both on BN-PAGE and SDS-PAGE (**Figure 2B**). Similarly, co-expression of heavy NRC2^EEE^-mCherry-4xMyc with light Rx-6xHA does not result in a size-shift for Rx in BN-PAGE relative to light NRC2^EEE^-4xMyc co-expressed with light Rx-6xHA. Again, a clear shift in size was observed in both the inactive and activated states for heavy NRC2^EEE^-mCherry-4xMyc relative to light NRC2^EEE^-4xMyc both in BN-PAGE and SDS-PAGE (**Figure 2B**). All in all, these data suggest that Rx and NRC2 likely do not form stable complexes with each other at resting state, and that following activation by its upstream sensor, NRC2 oligomerizes and forms a higher order complex that does not include Rx.

**Figure 2:**
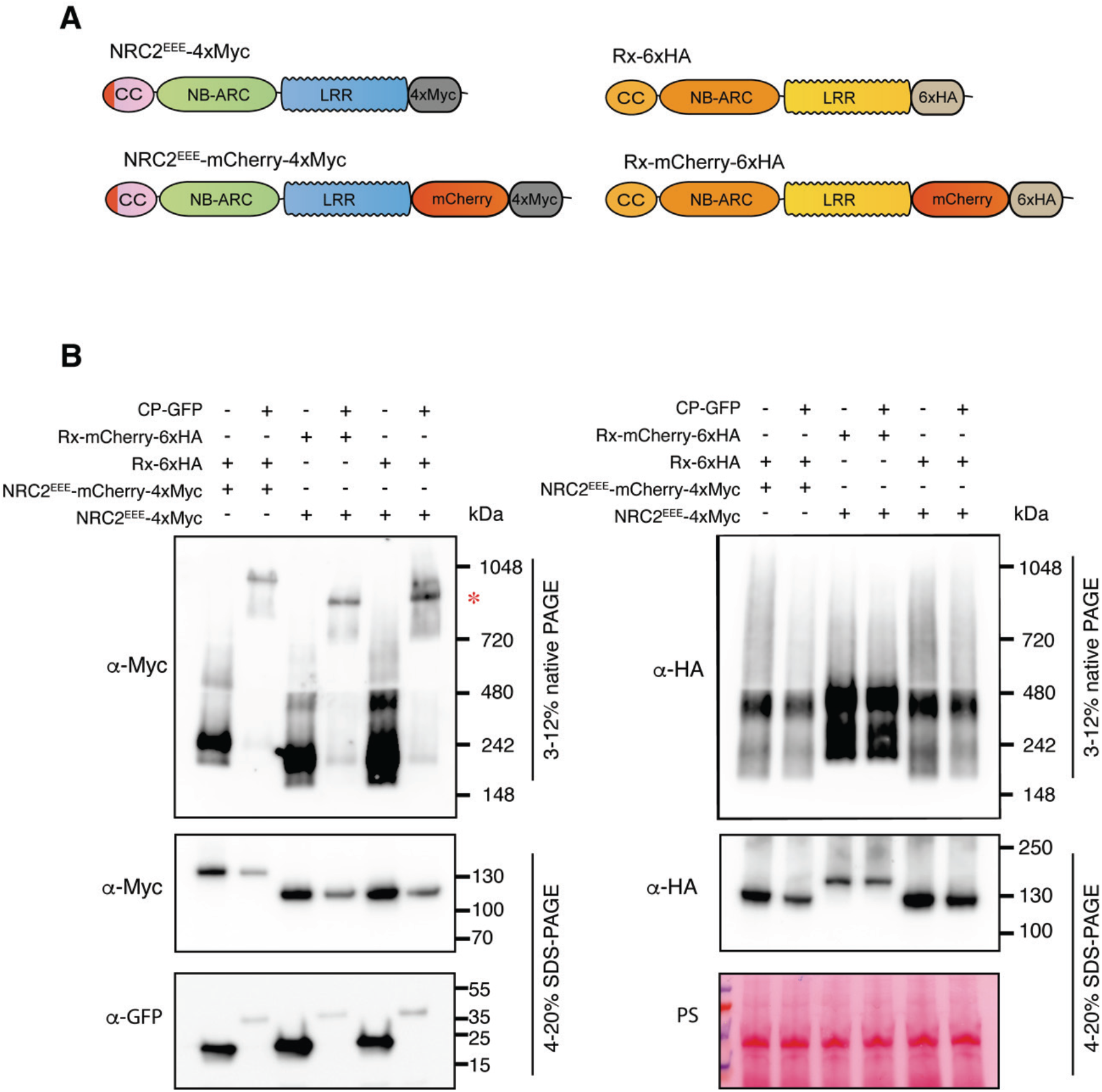
Rx and NRC2 do not form a stable complex upon activation. (**A**) Schematic representation of NRC2^EEE^ and Rx with differently sized tags. The C-terminally 4xMyc-tagged NRC2^EEE^ and C-terminally 6xHA tagged Rx used in previous experiments were termed “light” versions, while the new C-terminally tandem mCherry-4xMyc-tagged NRC2^EEE^ and C-terminally mCherry-6xHA-tagged Rx variants were termed “heavy” versions. (**B**) BN-PAGE and SDS-PAGE assays with inactive and activated Rx-NRC2 using different heavy and light sensor-helper combinations. Total protein extracts were run on native and denaturing PAGE assays in parallel and immunoblotted with the appropriate antisera labelled on the left. Approximate molecular weights (kDa) of the proteins are shown on the right. Red asterisk indicates bands corresponding to the activated NRC2 complex. Rubisco loading control was carried out using Ponceau stain (PS). The experiment was repeated three times.

### The activated NRC2 complex is composed of multiple NRC2 proteins

Considering that our findings are in line with the hypothesis that Rx does not form part of the active NRC2 complex, we next sought to further characterize the nature of this high molecular weight NRC2 complex. To do so, we leveraged the previously used heavy and light tag approach. We postulated that if NRC2 monomers were indeed forming a complex upon activation, a heterogeneous mixture of differently sized NRC2 molecules would lead to a shift in size of the activated complex, relative to homogeneous heavy or light NRC2 complexes. To test this hypothesis, we co-expressed Rx-6xHA with either light NRC2^EEE^-3xFLAG (104 kDa), heavy NRC2^EEE^-mCherry-4xMyc (133 kDa) or a mixture of heavy and light NRC2 variants in leaves of *nrc2/3/4 N. benthamiana* CRISPR mutant lines (**Figure 3A**). As expected, both the inactive and activated heavy NRC2^EEE^-mCherry-4xMyc exhibited a higher molecular weight than the inactive and activated light NRC2^EEE^-3xFLAG complexes, respectively (**Figure 3B)**. However, upon co-expressing both heavy NRC2^EEE^-mCherry-4xMyc and light NRC2^EEE^-3xFLAG, the activated NRC2 complex was of intermediate molecular weight, relative to either light NRC2^EEE^-3xFLAG or heavy NRC2-mCherry-4xMyc complexes (**Figure 3B**). The observation that combining differently sized NRC2 variants results in an intermediate molecular weight activated complex suggests that upon activation by Rx, both NRC2 versions are entering a complex. Interestingly, we were unable to see any change in size for inactive NRC2^EEE^ when mixing the two molecular weight variants (**Figure 3B**). We conclude that the fast-migrating lower molecular weight band observed for inactive NRC2 is likely not a complex of multiple NRC2 molecules. This allows us to draw a model for Rx-NRC2 activation in which, following effector-triggered activation, Rx leads to activation of its downstream helper NRC2, which oligomerizes and forms a resistosome complex that includes multiple NRC2 units.

**Figure 3:**
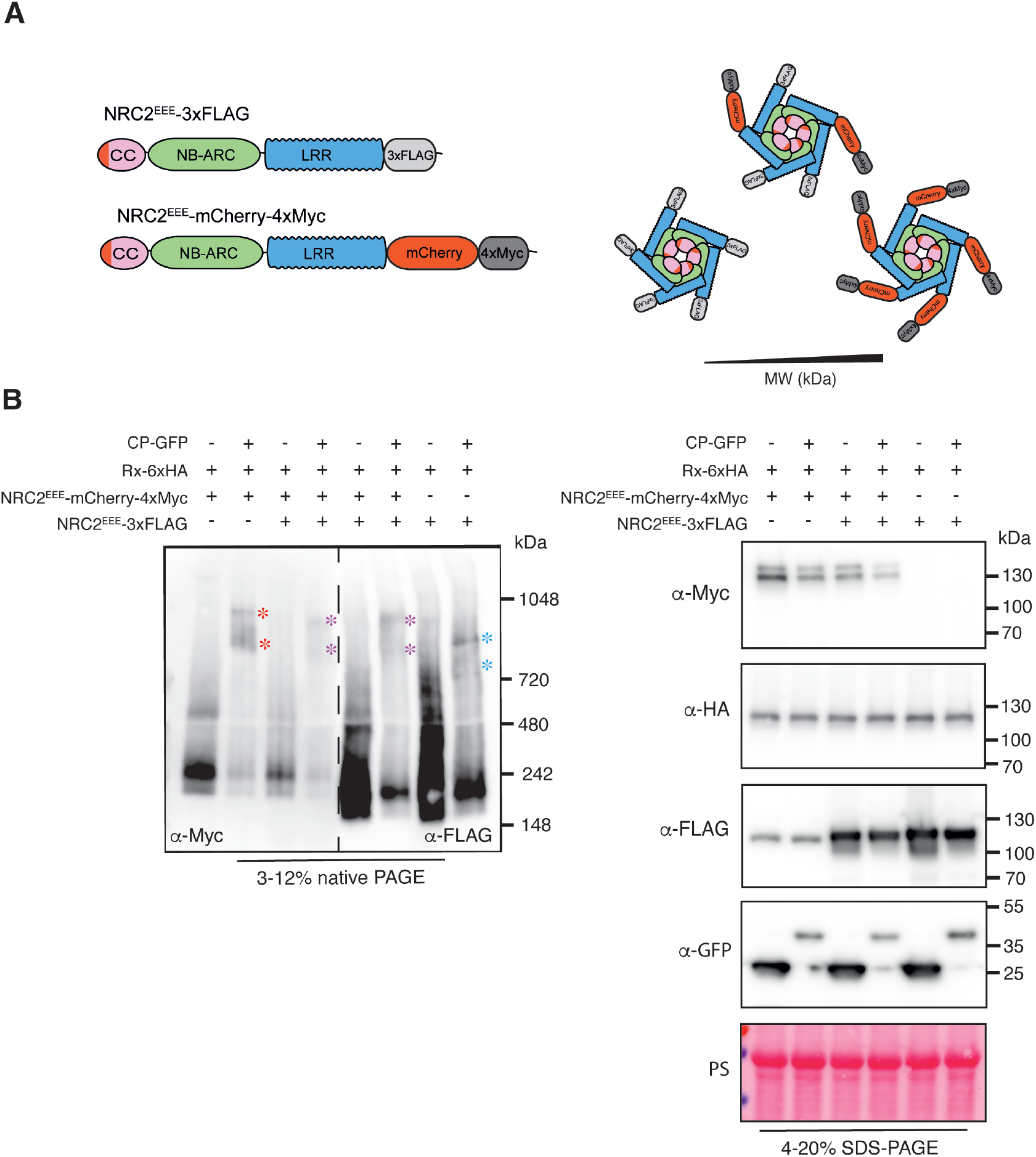
The activated NRC2^EEE^ complex is composed of multiple NRC2 units. (**A**) Schematic representation of NRC2^EEE^ with differently sized tags. The C-terminally 3xFLAG-tagged NRC2^EEE^ was termed the “light” version and C-terminally mCherry-4xMyc tagged NRC2EEE was termed the “heavy” variant. If NRC2 forms homo-oligomers, active complexes with different proportions of “heavy” and “light” NRC2 monomers should in theory exhibit different molecular weights. (**B**) BN-PAGE and SDS-PAGE assays with inactive and activated Rx-NRC2 using different “heavy” and “light” helper combinations. Total protein extracts were run on native and denaturing PAGE assays in parallel and immunoblotted with the appropriate antisera labelled on the left. Approximate molecular weights (kDa) of the proteins are shown on the right. A non-specific band was observed at ∼110 kDa for the anti-FLAG antibody. Red asterisks indicate bands corresponding to the activated “heavy” NRC2 complexes. Blue asterisks indicate bands corresponding to “light” NRC2 complexes. Purple bands indicate intermediate molecular weight complexes combining “heavy” and “light” NRC2. Rubisco loading control was carried out using Ponceau stain (PS). The experiment was repeated three times.

### Bs2, another NRC2-dependent sensor NLR, also triggers oligomerization of NRC2

Since Rx is able to trigger oligomerization of the downstream helper NRC2, we tested whether this applies to other NRC-dependent sensors. We performed complementation assays co-expressing NRC2^EEE^ in leaves of *nrc2/3/4 N. benthamiana* CRISPR mutants along with the inactive or activated NRC2/3/4-dependent sensor Bs2, or Rpi-blb2, an NRC4-dependent sensor that is unable to signal through NRC2 (**Figure 4A**) (Duggan *et al*., 2021; Wu *et al*., 2017). While Bs2 activation with AvrBs2 was able to trigger NRC2^EEE^ resistosome formation in a similar fashion to Rx, Rpi-blb2 activation with AVRblb2 did not lead to the formation of a NRC2^EEE^ resistosome (**Figure 4B**). This suggests that sensor-mediated NRC oligomerization is a general principle for NRC dependent sensors. Moreover, the observation that only sensors that genetically require NRC2 can trigger its oligomerization shows that previously genetically characterized sensor-helper specificities in the NRC network can be recapitulated biochemically by helper oligomerization in BN-PAGE.

**Figure 4:**
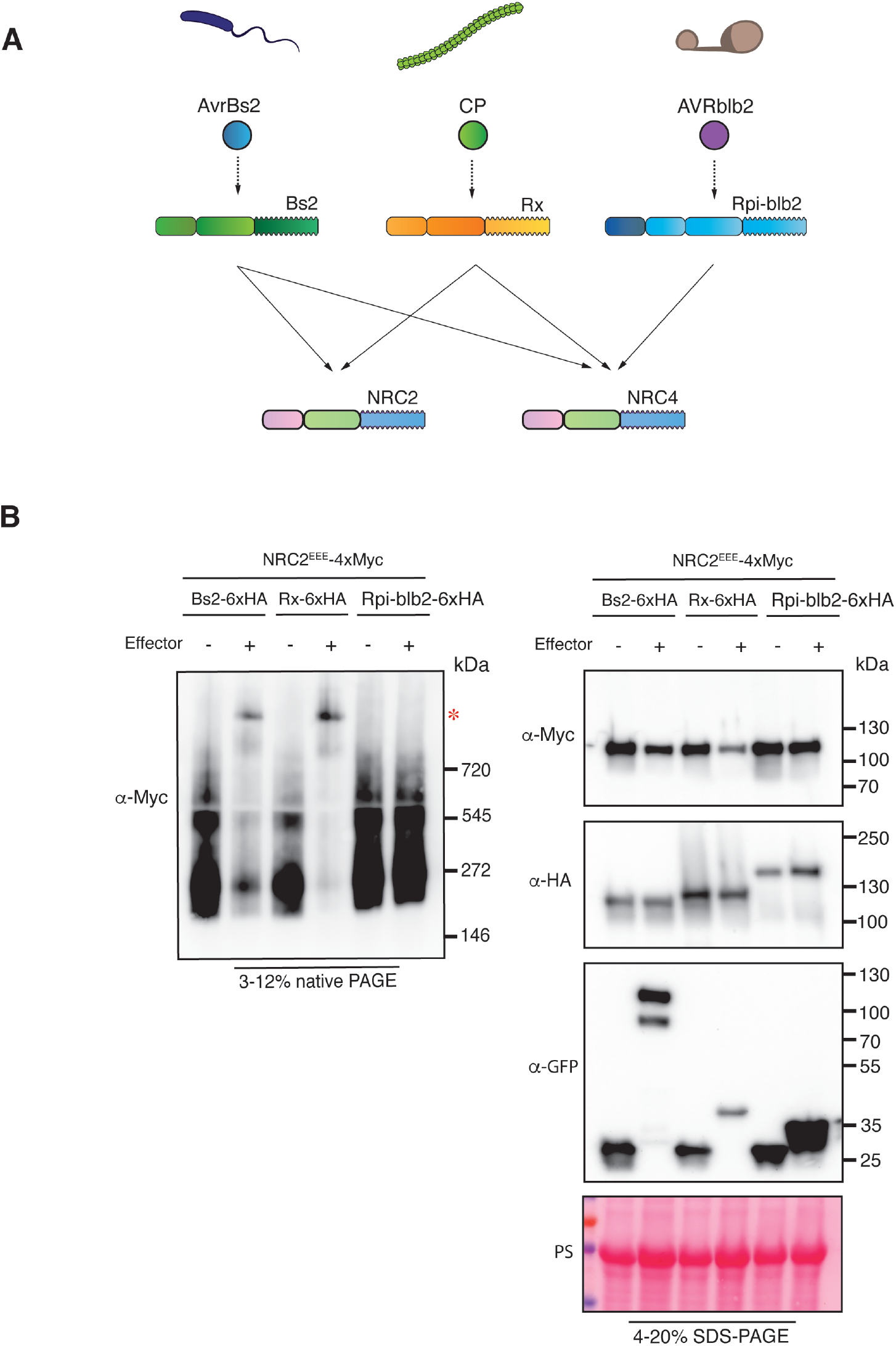
Bs2, another NRC2-dependent sensor NLR, triggers oligomerization of NRC2^EEE^. (**A**) Schematic representation of the NRC genetic dependencies of sensors used in this experiment. (**B**) BN-PAGE and SDS-PAGE assays with inactive and activated NRC-dependent sensors and NRC2. Total protein extracts were run on native and denaturing PAGE assays in parallel and immunoblotted with the appropriate antisera labelled on the left. Approximate molecular weights (kDa) of the proteins are shown on the right. Red asterisk indicates bands corresponding to the activated NRC2 complexes. Rubisco loading control was carried out using Ponceau stain (PS). The experiment was repeated three times.

### Activated NRC2 oligomers accumulate in membrane-associated puncta whereas Rx remains cytoplasmic

The cellular biology of activated NLRs remains poorly understood. We previously showed that the NRC helper NLR NRC4 exhibits dynamic spatiotemporal changes in subcellular localization following effector-triggered activation of its upstream sensor NLR Rpi-blb2 (Duggan *et al*., 2021). We applied similar methods to study the sub-cellular dynamics of the Rx-NRC2 system by transiently co-expressing fluorescently tagged versions of Rx-RFP and NRC2^EEE^-GFP in leaves of *nrc2/3/4 N. benthamiana* CRISPR mutants. We activated the Rx-NRC2 system by expressing 4xMyc-tagged CP or a 4xMyc-tag empty vector (EV) control and monitored sensor and helper localization using confocal live-cell imaging. As a plasma membrane (PM) marker, we co-expressed RPW8.2-BFP (Duggan *et al*., 2021). In parallel, protein was extracted from the same leaf tissue used for microscopy to confirm that the tags do not interfere with Rx-mediated NRC2 cell death and oligomerization by BN-PAGE assays (**Figure S3**). In their inactive state, both Rx-RFP and NRC2^EEE^-GFP co-localize to the cytoplasm in 100% of observations (N = 16 images) (**Figure 5A, Movie S1**). Strikingly, when co-expressing CP, activated NRC2^EEE^-GFP predominantly localizes to puncta which frequently co-localize with the PM, marked by RPW8.2-BFP. In contrast, Rx-RFP does not exhibit major changes in subcellular localization. The sensor remains in the cytoplasm and does not concentrate in the NRC2 puncta (15/16 images taken). These puncta are uniformly distributed throughout the PM (**Figure 5B, Movie S1**). To investigate the membrane association of the activated oligomeric NRC2 complex, we obtained protein extracts from the same tissues used for microscopy and carried out membrane fractionation assays in nondenaturing conditions using the same experimental setup described above and performed SDS-PAGE assays using the different fractions (**Figure 5C**). In line with what we previously observed in live-cell imaging experiments, we found that in the inactive state, both Rx and NRC2^EEE^ are mainly present in the soluble fraction. Following activation, NRC2^EEE^ is equally distributed between the soluble and membrane fractions, indicating a shift in subcellular localization and increased membrane-association. Rx, however, exhibits no such shift upon activation and remains predominantly in the soluble fraction. We conclude that upon effector-triggered activation of Rx, the sensor subsequently mediates activation of its helper NRC2^EEE^ in the cytoplasm. The activated NRC2 units form oligomeric resistosomes that dynamically re-localize and form membrane-associated puncta that are separate from the sensor.

**Figure 5:**
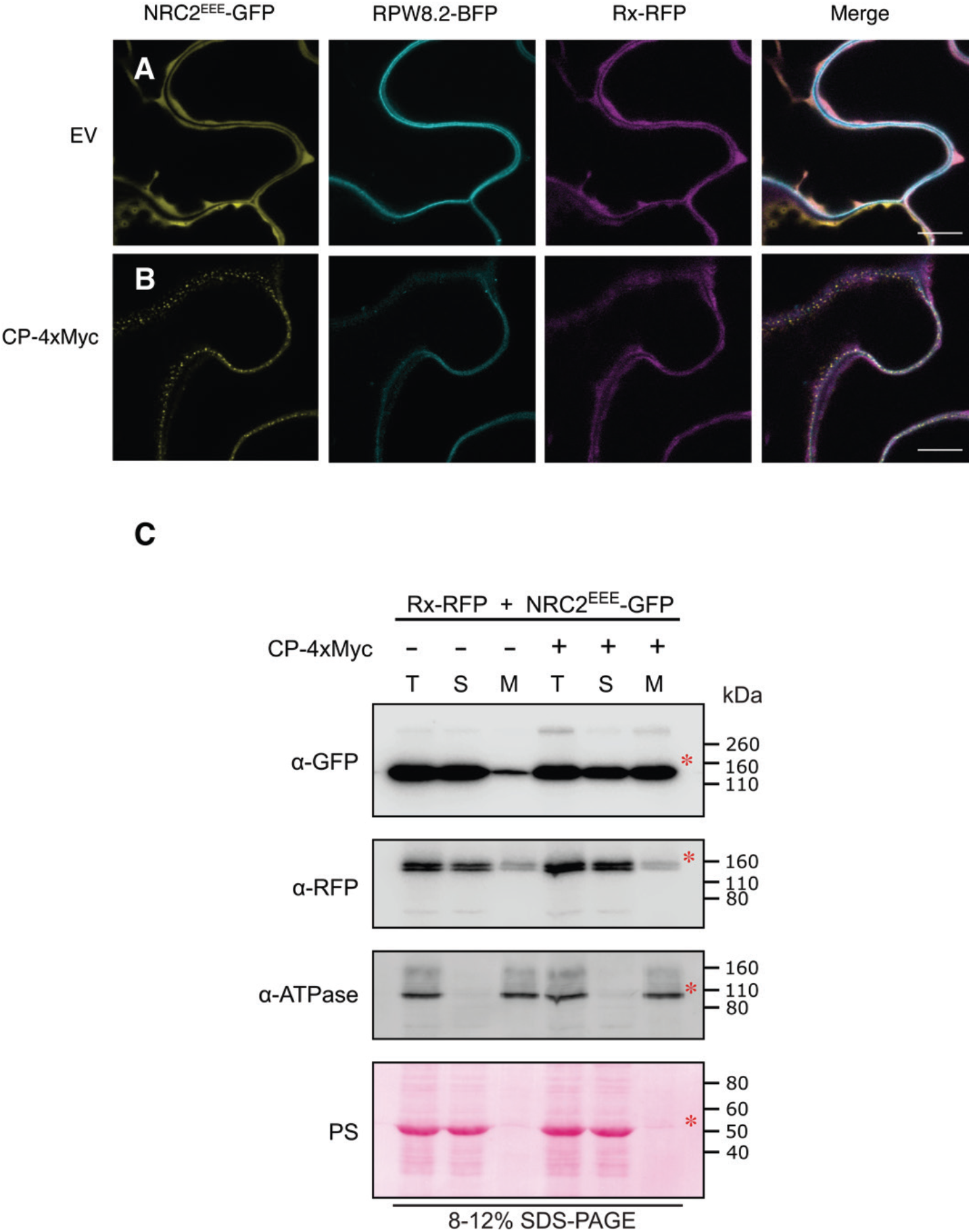
*Potato virus X* coat protein activated NRC2 forms plasma membrane-associated puncta, unlike Rx. C-terminally GFP-tagged NRC2^EEE^ and C-terminally RFP-tagged Rx were co-expressed either with an EV-4xMyc construct or a CP-4xMyc construct in leaves of *nrc2/3/4 N. benthamiana* CRISPR mutant lines. (**A**-**B**) Single-plane confocal micrographs show the localization of both components of the inactive and active Rx-NRC2 system, together with PM marker RPW8.2-BFP. Scale bars represent 10 μm. (**A**) NRC2^EEE^-GFP and Rx-RFP co-localize in the cytoplasm prior to activation. (**B**) Upon co-expression of CP and activation of the system, NRC2^EEE^ forms puncta associated with the PM while Rx remains in the cytoplasm. (**C**) Membrane enrichment assays are consistent with microscopy, showing that inactive NRC2^EEE^-GFP is mostly present in the soluble (cytoplasmic) fraction, whereas activated NRC2^EEE^-GFP exhibits equal distribution across soluble and membrane fractions. Rx is mostly present in the soluble fraction and exhibits no change upon activation of the system with CP. T = total, S = soluble, M = membrane. ATPase was used as a membrane marker. Rubisco was used as a marker for total and soluble fractions and visualized by Ponceau staining (PS). Red asterisks indicate bands matching the expected MW for each protein. The experiment was repeated two times.

### *Potato virus X* infection triggers Rx-mediated oligomerization of NRC2

To confirm our previous results in the context of pathogen infection rather than with activation using heterologously expressed effectors, we used BN-PAGE to study the activation of NRC2 in the context of viral infection. We transiently co-expressed Rx and NRC2^EEE^ in leaves of *nrc2/3/4 N. benthamiana* CRISPR mutant lines and activated the system by infecting leaf tissues with PVX via co-expressing a GFP tagged PVX variant (pGR106::PVX::GFP), as described previously (Derevnina *et al*., 2021). Transient expression of free GFP was used as a negative control for infection. After 3 days, protein extracts from infected or uninfected leaf tissues were used for BN-PAGE assays (**Figure 6A**). Consistent with previous results, infection with PVX led to sensor-dependent oligomerization of NRC2^EEE^ (**Figure 6B**). Interestingly, in the SDS-PAGE, a very strong GFP signal was observed in all PVX treatments, suggesting that the virus is able to replicate in the presence of Rx and NRC2^EEE^. This implies that, while the leucine to glutamic acid mutations in the N-terminal MADA motif of NRC2 do not prevent resistosome formation mediated by Rx, the NRC2^EEE^ mutant is both unable to trigger hypersensitive cell death and mediate Rx-dependent PVX resistance (**Figure 6B**). We conclude that for sensor-helper pairs in the NRC network, the sensor NLR is activated following pathogen recognition and subsequently signals to its downstream NRC helper, leading to its oligomerization and resistosome formation. We also conclude that an NRC helper with a functional MADA-motif is necessary for the Rx-NRC2 system to mediate full PVX resistance.

**Figure 6:**
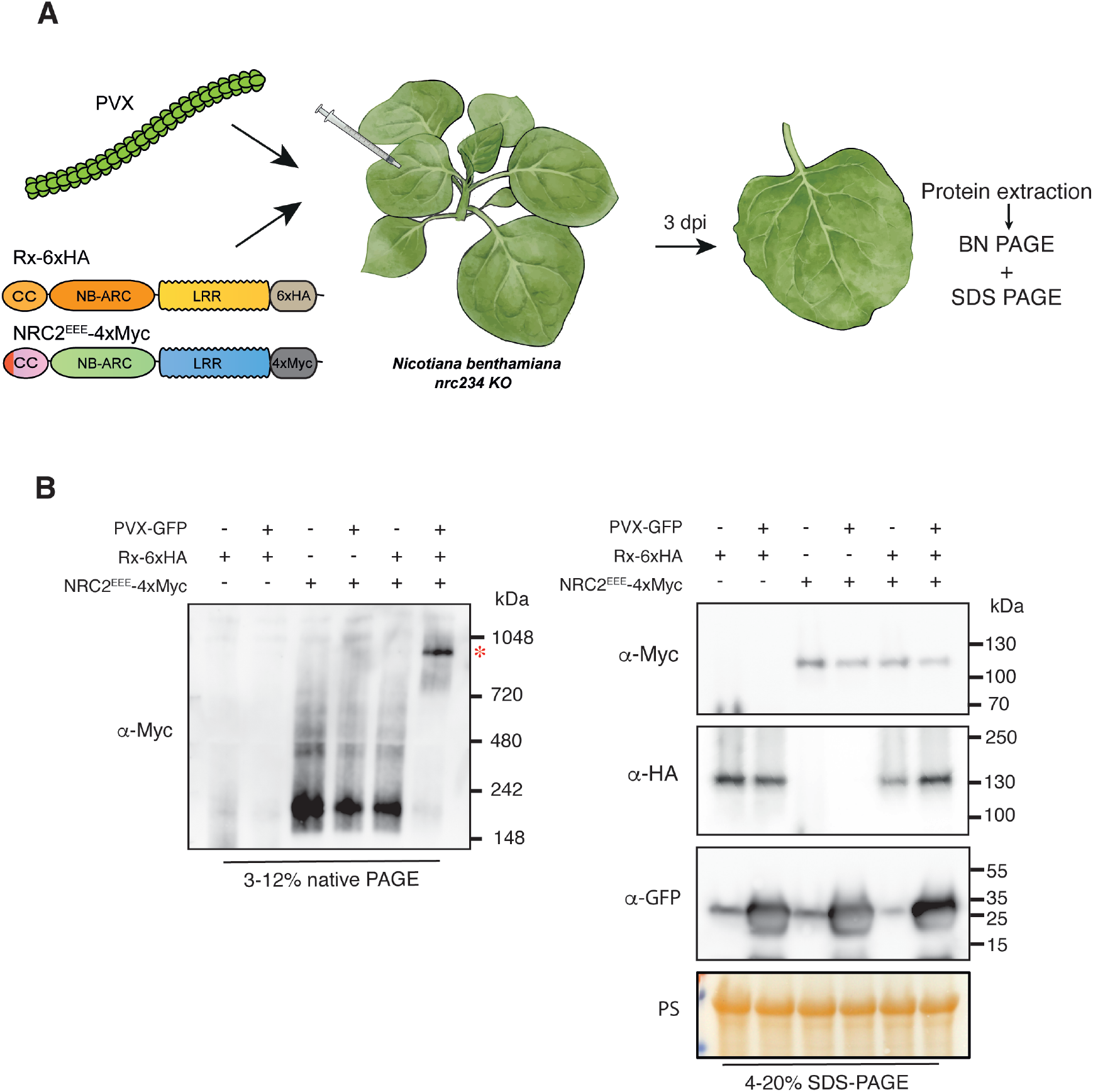
*Potato virus X* infection triggers Rx-mediated oligomerization of NRC2. (**A**) Schematic representation of the experimental pipeline used. *A. tumefaciens* strains were used to transiently express proteins of interest in leaves of *nrc/2/3/4 N. benthamiana* CRISPR mutant lines by agroinfiltration. Simultaneously, the same leaves were infected with PVX by using an *A. tumefaciens* carrying GFP expressing PVX. Free GFP was used as a negative control for PVX-GFP infection. Free mCherry-4xMyc and mCherry-6xHA fusions were used as controls in the treatments without NRC2 and Rx, respectively. Leaf tissue was harvested 3 days post-infiltration and total protein extracts were used for BN and SDS-PAGE assays. (**B**) BN-PAGE and SDS-PAGE assays with infected and uninfected leaves expressing Rx and NRC2. Total protein extracts were run on native and denaturing PAGE assays in parallel and immunoblotted with the appropriate antisera labelled on the left. Approximate molecular weights (kDa) of the proteins are shown on the right. Red asterisks indicate bands corresponding to the activated NRC2 complexes. Rubisco loading control was carried out using Ponceau stain (PS). The experiment was repeated three times.

### *Potato virus X* infection triggers Rx-dependent formation of membrane-associated NRC2 puncta

We next sought out to study the sensor-dependent subcellular reorganization of NRC2 described above in the context of pathogen infection. To this end, we transiently co-expressed NRC2^EEE^ either with Rx-RFP or free RFP in leaves of *nrc2/3/4 N. benthamiana* CRISPR mutant lines and activated the system by infecting leaf tissues with PVX (pGR106::PVX), as described previously (Derevnina *et al*., 2021). After 3 days, we monitored sensor and helper localization using confocal live-cell imaging (**Figure 7**). Under these infection conditions, the helper NRC2^EEE^-GFP remains in the cytoplasm in the absence of its upstream sensor Rx (18/18 images taken) (**Figure 7A**). Consistent with previous results, when co-expressing NRC2^EEE^-GFP together with Rx-RFP and infecting with PVX, the helper predominantly localizes to PM-associated puncta. In contrast, Rx remains in the cytoplasm during PVX infection and does not exhibit co-localization with NRC2 (18/18 images taken) (**Figure 7B**). Our data indicate that the dynamic re-localization and PM-association of NRC2 we observed following treatment with CP (**Figure 6**) also occurs during pathogen infection.

**Figure 7:**
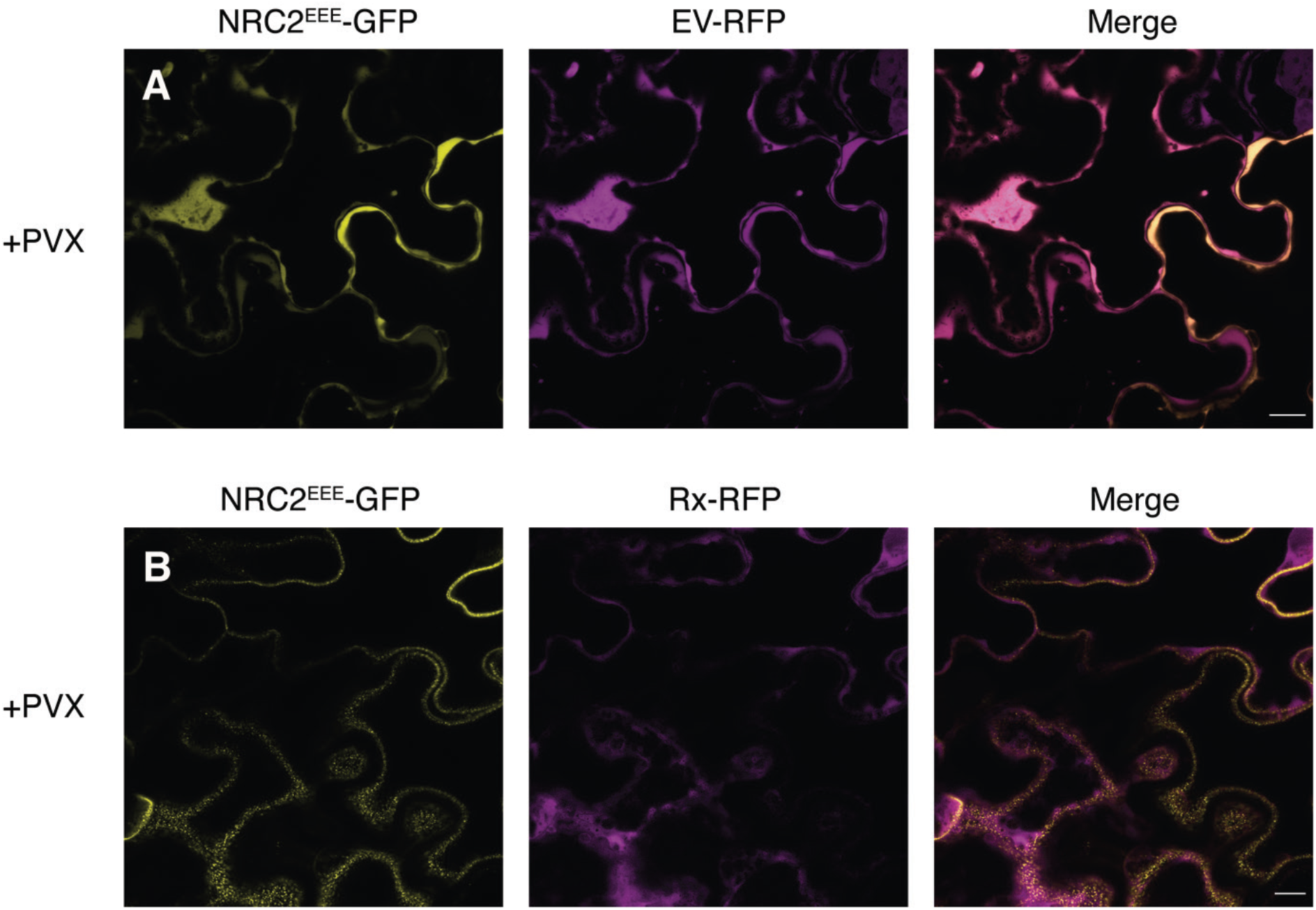
*Potato virus X* infection triggers Rx-dependent NRC2 membrane-associated puncta. (**A**-**B**) Single-plane confocal micrographs show the localization of NRC2 together with Rx or free RFP during PVX infection. C-terminally GFP-tagged NRC2^EEE^ was co-infiltrated with either EV-RFP or Rx-RFP in leaves of *nrc2/3/4 N. benthamiana* CRISPR mutant lines and infected with PVX by agroinfection. Scale bars represent 10 μm. (**A**) In the absence of Rx, NRC2^EEE^-GFP is localized to the cytoplasm during PVX infection. (**B**) When Rx is present, NRC2^EEE^ forms puncta associated with the PM during PVX infection.

## Discussion

The aim of this study was to develop a better understanding of the molecular mechanisms that underpin activation of paired and networked plant NLR immune receptors. Such *in planta* studies were previously hampered by the cell death response associated with activated NLRs. The capacity to make mutations in the N-terminal MADA motif of CC-NLR helper proteins that abolish elicitation of cell death without affecting activation enabled us to develop readouts for the formation of resistosome-like oligomers and address several questions related to NRC activation. We used biochemical and cellular biology techniques to develop a working model for activation of the NRC network sensor NLRs Rx and Bs2 and their helper NRC2 (Derevnina *et al*., 2021; Wu *et al*., 2017). Our data demonstrate that these sensor NLRs can mediate NRC2 oligomerization (**Figure 1, Figure 4**) upon activation with their corresponding effectors and in the case of Rx, during pathogen infection (**Figure 6**). We show that sensor activation also mediates helper re-localization to PM during pathogen infection (**Figure 5, Figure 7**). This activated NRC2 complex appears to be an oligomer containing multiple NRC2 units, which excludes the sensor NLR (**Figure 1, Figure 2, Figure 3**), suggesting a model for sensor-helper activation that differs from mammalian paired NLR systems, such as NAIP/NLRC4 (Vance, 2015; Zhang *et al*., 2015). These findings and those of a co-published study on oligomerization of NRC2 after activation of the oomycete resistance proteins Rpi-amr1 and Rpi-amr3 (Ahn *et al*., 2022) have led us to develop an activation-and-release model for NLRs in the NRC network (**Figure 8**).

**Figure 8:**
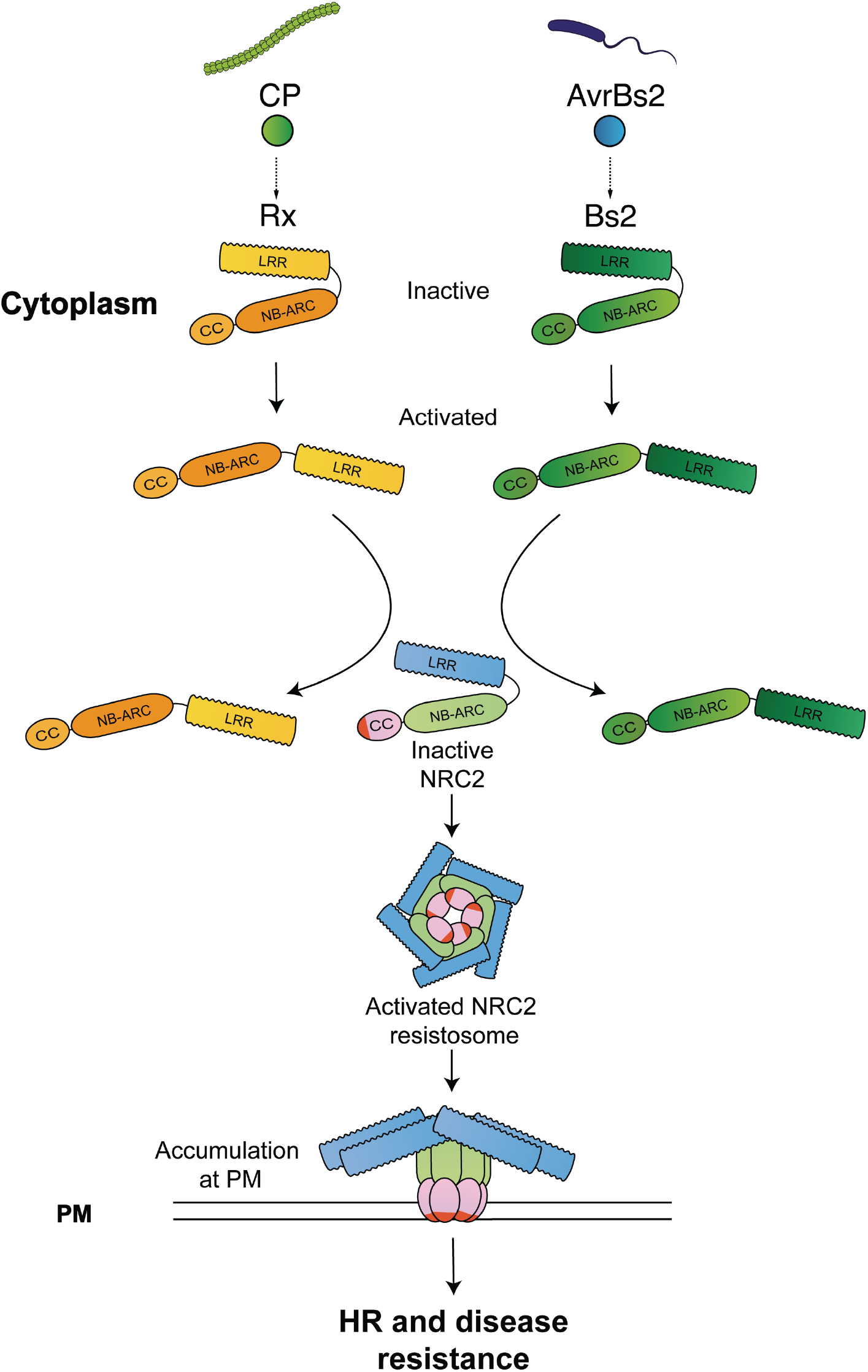
An activation-and-release working model for sensor-helper pairs in the NRC network. Prior to effector-triggered activation, NRC-dependent sensors such as Rx and Bs2 are held in an inactive conformation by intramolecular interactions. Upon perceiving their cognate effectors, the sensors undergo a series of conformational changes that allow them to signal to NRC2 and mediate its homo-oligomerization and resistosome formation. The activated NRC2 resistosome separates from the sensors and accumulates at the PM. The sensors remain in the cytoplasm, separate to the activated helper.

The N-terminal α1-helices of ZAR1, Sr35 and the NRCs are defined by a molecular signature called the “MADA motif” (Adachi *et al*., 2019a; Kourelis *et al*., 2021a). This motif can be found in ∼20% of angiosperm CC-NLRs and is essential for immune signalling and disease resistance. For TIR-NLR biology, multiple essential downstream components for immune signaling, such as the EDS1 signaling hub and the CC_R_ NLRs NRG1 and ADR1 have been identified, which enabled the generation of genetic backgrounds with which to study TIR-NLR activities in the absence of immune signaling (Gantner *et al*, 2019; Lapin *et al*, 2019; Martin *et al*., 2020; Saile *et al*, 2020; Sun *et al*, 2021). Unlike TIR-NLRs, approaches to study activated CC-NLRs in the absence of a plant cell death response were not available until recently (Adachi *et al*., 2019a; Duggan *et al*., 2021). The availability of MADA mutant variants that no longer trigger cell death, but retain the capacity to activate, oligomerize and re-localize enabled us to study the cellular and biochemical mechanisms of CC-NLR activation.

MADA motif mutants of NRC2 are unable to mediate cell death yet they can still oligomerize and relocalize to the PM upon activation by their upstream sensors. How does this fit with current models of CC-NLR execution of the hypersensitive cell death? Our work complements existing studies that show how MADA mutants can be leveraged to shed light on the diversity of molecular mechanisms that underpin CC-NLR activation (Adachi *et al*., 2019a; Förderer *et al*., 2022; Hu *et al*., 2020; Kourelis *et al*., 2021a). Two studies on NRC4 and ZAR1 respectively, showed that mutations in the N-terminal MADA-motif containing α1-helices abolished cell death triggered by these NLRs but had no effect on PM localization (Duggan *et al*., 2021; Hu *et al*., 2020). Similar to NRC2^EEE^, ZAR1 and Sr35 with mutated N-termini were still able to form resistosomes (Förderer *et al*., 2022; Hu *et al*., 2020). Resistosome assembly and PM localization are, therefore, required but not sufficient to mediate cell death. In the case of Sr35, mutations in the MADA motif even abolished the capacity of this NLR to trigger cell death in insect cells (Förderer *et al*., 2022), suggesting that no further downstream components are required or that the signalling initiated by these resistosomes can engage with highly conserved pathways present across the plant and animal kingdoms. How do MADA motif mutations interfere with cell death? One possibility is that the MADA mutations interfere with the Ca^2+^ channel activity that was recently assigned to activated CC and CC_R_-NLR proteins (Bi *et al*., 2021; Förderer *et al*., 2022; Jacob *et al*., 2021). Alternatively,

N-terminal α1-helices in CC-NLRs with mutated MADA motifs may result in resistosomes that associate with the plasma membrane but are unable to fully penetrate the lipid bilayer to form a functional pore or channel. Further research will dissect the precise role of this N-terminal motif in NRC and CC-NLR-mediated cell death.

The recent elucidation of multiple plant NLR structures has demonstrated that plant and metazoan NLRs exhibit functional differences despite several commonalities. In this work, we propose an activation-and-release model for sensor-helper pairs in the NRC network (**Figure 8**). In this working model, effector-triggered activation of a sensor NLR leads to conformational changes, which relay a signal to downstream helpers, potentially via transient interactions. These helpers subsequently activate, oligomerize and form resistosomes. The activated helper complexes then part ways with their sensors and re-localize to the plasma membrane where they trigger cell death. It is possible that transient intermediates exist where sensors interact with their helpers to trigger their activation. However, BN-PAGE assays with differently sized versions of Rx and NRC2 and confocal microscopy suggest that a stable hetero-complex scenario is unlikely for the activated Rx- NRC2 system. This points to a biochemical model for plant paired NLR activation that differs from activation processes previously characterized for metazoan NLR pairs. We conclude that these plant and metazoan paired and networked NLRs exhibit distinct activation mechanisms and biochemical processes. Whether this activation-and-release model applies to other paired plant CC-NLRs or even other paired metazoan NLRs remains to be tested.

Previous work has shown that following CP-triggered activation, Rx undergoes a series of conformational changes that lead to cell death and immune activation (Moffett *et al*., 2002; Rairdan & Moffett, 2006). Nonetheless, how a signal is relayed from sensor to helper remains unknown. While Rx does not oligomerize upon activation, the conformational switch may allow Rx to interact transiently with NRC2 in order to mediate its activation. To date, conclusive evidence that NRC-dependent sensors and their NRC helpers form stable complexes has not been obtained, possibly because the complexes are transient. Regardless of whether a direct or indirect interaction between sensor and helper mediates NRC activation, these findings and those of the companion study (Ahn *et al*., 2022) indicate that the mature NRC2 resistosome is released from the activated sensor. In this scenario, one Rx or Bs2 could potentially activate multiple NRC2 molecules, possibly triggering an NRC2 oligomerization cascade independent of the sensor. Alternatively, NRC2 may form a transient sensor-helper heterocomplex with the sensor, which could act as an intermediate polymerization scaffold for the NRC2 resistosome, reminiscent of the first stages of NAIP/NLRC4 inflammasome maturation. A mechanism in which one sensor molecule can activate multiple NRC resistosomes would be much more efficient in amplifying immune signals as opposed to an activated sensor stably engaging in a sensor-helper heterocomplex. Such an amplification would be analogous to the working model for TIR-NLR/CC_R_-NLR sensor-helper pairs, where small molecules produced by activated TIR-NLR sensors lead to downstream helper activation via the EDS1 signaling hub, triggering CC_R_-NLR resistosome formation (Huang *et al*, 2022; Jacob *et al*., 2021; Sun *et al*., 2021). Further studies will hopefully shed more light on the dynamics of NRC resistosome assembly.

The PVX system allowed us to study paired NLR activation during pathogen infection, taking the state-of-the-art beyond activation with effector proteins (**Figure 6, Figure 7**). This work complements previous studies on NLR oligomerization upon heterologous expression of cognate effectors (Duxbury *et al*, 2020; Hu *et al*., 2020; Li *et al*., 2020; Ma *et al*., 2020; Martin *et al*., 2020; Williams *et al*, 2014), showing that the same mechanism likely applies during infection by pathogenic organisms. Investigating the oligomeric state and subcellular localization of paired/networked NLRs upon infection will provide further insights into the mechanisms that underpin NLR-mediated immunity. In the case of NRC4, this helper can focally accumulate at the interface between *P. infestans* and the host plant at the site where effectors are delivered before re-localizing and forming discrete puncta at the PM following activation (Duggan *et al*., 2021). Interestingly, the puncta observed for activated NRC2 and NRC4 are distributed throughout the cell PM. What is the exact nature of these puncta, how many resistosomes accumulate in the observed PM micro-domains and whether they form macro-complexes remain open questions.

A multitude of structural, biochemical and cell biology studies have contributed to our mechanistic understanding of NLR activation and signalling (Duxbury *et al*., 2021; Kourelis & Van Der Hoorn, 2018; Ngou *et al*., 2022). Our work expands on the current conceptual mechanistic framework of NLR activation by incorporating the higher order of genetic complexity presented by plant immune receptor networks (Wu *et al*., 2017; Wu *et al*., 2018). In the case of singleton NLRs, an individual protein can perceive effectors and subsequently form a resistosome to execute cell death. We previously proposed that in plant NLR networks, sensors have specialized over evolutionary time to perceive effectors and have lost the biochemical features required for executing cell death (Adachi *et al*., 2019a; Adachi *et al*., 2019b). Based on our current working model (Figure 8), sensor NLRs have lost the capacity to oligomerize into resistosomes and have specialized in pathogen perception and helper activation. In contrast, activated NRC2 behaves similarly to other MADA-CC-NLRs by forming resistosome-like homo-oligomers that translocate to the PM. Nonetheless, the many-to-many sensor-helper connections in the NRC network raise further questions. What is the precise nature of the activation signal relayed from sensor to helper? What are the precise dynamics of NRC resistosome assembly? How do the molecular determinants for sensor-helper specificity translate into resistosome formation? Addressing these questions holds the potential to advance our understanding of the diversity of plant NLR immune activation beyond functional singleton NLRs.

## Materials and methods

### Plant growth conditions

Wild-type and *nrc2/3/4* CRISPR mutant *N. benthamiana* lines were grown in a controlled environment growth chamber with a temperature range of 22 to 25 ºC, humidity of 45% to 65% and a 16/8-hour light/dark cycle.

### Plasmid constructions

The Golden Gate Modular Cloning (MoClo) kit (Weber *et al*, 2011) and the MoClo plant parts kit (Engler *et al*, 2014) were used for cloning, and all vectors are from this kit unless specified otherwise. Effectors, receptors and NRCs were cloned into the binary vector pJK268c, which contains the Tomato bushy stunt virus silencing inhibitor p19 in the backbone (Kourelis *et al*, 2020). Cloning design and sequence analysis were done using Geneious Prime (v2021.2.2; https://www.geneious.com). Plasmid construction is described in **Table S1**.

### Cell death assays by agroinfiltration

Effectors and NLR immune receptors of interest were transiently expressed according to previously described methods (Bos *et al*, 2006). Briefly, leaves from 4–5-week-old plants were infiltrated with suspensions of *A. tumefaciens* GV3101 pM90 strains transformed with expression vectors coding for different proteins indicated. Final OD_600_ of all *A. tumefaciens* suspensions were adjusted in infiltration buffer (10 mM MES, 10 mM MgCl_2,_ and 150 μM acetosyringone (pH 5.6)). Final OD_600_ used was 0.3 for each NLR immune receptor and 0.2 for all effectors, adding up to a total OD_600_ of 0.8.

### Extraction of total proteins for BN-PAGE and SDS-PAGE assays

Four to five-week-old plants were agroinfiltrated as described above with constructs of interest and leaf tissue was collected 3 days post agroinfiltration. Final OD_600_ used was 0.3 for each NLR immune receptor used and 0.2 for all effectors for a total OD_600_ of 0.8. BN-PAGE was performed using the Bis-Tris Native PAGE system (Invitrogen) according to the manufacturer’s instructions. Leaf tissue was ground using a Geno/Grinder tissue homogenizer and total protein was subsequently extracted and homogenized in GTMN extraction buffer (10% glycerol, 50 mM Tris-HCl (pH 7.5), 5 mM MgCl_2_ and 50 mM NaCl) supplemented with 10 mM DTT, 1x protease inhibitor cocktail (SIGMA) and 0.2% Nonidet P-40 Substitute (SIGMA). Samples were incubated in extraction buffer on ice for 10 minutes with short vortex mixing every 2 minutes. Following incubation, samples were centrifuged at 5,000 x*g* for 15 minutes and the supernatant was used for BN-PAGE and SDS-PAGE assays.

### BN-PAGE assays

For BN-PAGE, samples extracted as detailed above were diluted as per the manufacturer’s instructions by adding NativePAGE 5% G-250 sample additive, 4x Sample Buffer and water. After dilution, samples were loaded and run on Native PAGE 3%-12% Bis-Tris gels alongside either NativeMark unstained protein standard (Invitrogen) or SERVA Native Marker (SERVA). The proteins were then transferred to polyvinylidene difluoride membranes using NuPAGE Transfer Buffer using a Trans-Blot Turbo Transfer System (Bio-Rad) as per the manufacturer’s instructions. Proteins were fixed to the membranes by incubating with 8% acetic acid for 15 minutes, washed with water and left to dry. Membranes were subsequently re-activated with methanol in order to correctly visualize the unstained native protein marker. Membranes were immunoblotted as described below.

### SDS-PAGE assays

For SDS-PAGE, samples were diluted in SDS loading dye and denatured at 72 ºC for 10 minutes. Denatured samples were spun down at 5,000 x*g* for 3 minutes and supernatant was run on 4%-20% Bio-Rad 4%-20% Mini-PROTEAN TGX gels alongside a PageRuler Plus prestained protein ladder (Thermo Scientific). The proteins were then transferred to polyvinylidene difluoride membranes using Trans-Blot Turbo Transfer Buffer using a Trans-Blot Turbo Transfer System (Bio-Rad) as per the manufacturer’s instructions. Membranes were immunoblotted as described below.

### Immunoblotting and detection of BN-PAGE and SDS-PAGE assays

Membranes were blocked with 5% milk in Tris-buffered saline plus 0.01% Tween 20 (TBS-T) for an hour at room temperature and subsequently incubated with desired antibodies at 4 ºC overnight. Antibodies used were anti-GFP (B-2) HRP (Santa Cruz Biotechnology), anti-HA (3F10) HRP (Roche), anti-Myc (9E10) HRP (Roche), and anti-FLAG (M2) HRP (Sigma), all used in a 1:5000 dilution in 5% milk in TBS-T. To visualize proteins, we used Pierce ECL Western (32106, Thermo Fisher Scientific), supplementing with up to 50% SuperSignal West Femto Maximum Sensitivity Substrate (34095, Thermo Fishes Scientific) when necessary. Membrane imaging was carried out with an ImageQuant LAS 4000 or an ImageQuant 800 luminescent imager (GE Healthcare Life Sciences, Piscataway, NJ). Rubisco loading control was stained using Ponceau S (Sigma) or Ponceau 4R (AG Barr).

### BN and SDS-PAGE assays with PVX infection (agroinfection)

Four to five-week-old plants were agroinfiltrated as described above with constructs of interest. Simultaneously, PVX was delivered by agroinfection using an *A. tumefaciens* strain carrying GFP-labelled PVX (pGR106-PVX-GFP). Final OD_600_ used was 0.3 for each NLR immune receptor used and 0.05 for the *A. tumefaciens* strain carrying PVX or free GFP for a total OD_600_ of 0.65. Leaf tissue was collected 3 days post agroinfiltration. BN-PAGE and SDS-PAGE assays were carried out as described above.

### Confocal microscopy

Three to four-week-old plants were agroinfiltrated as described above with constructs of interest. PVX was delivered as before, or the coat protein CP-4xMyc or EV control at OD_600_ 0.1; Rx-RFP at OD_600_ of 0.25 and NRC2^EEE^-GFP at OD_600_ of 0.25. Leaf tissue was prepared for imaging by sectioning of desired area surrounding an infection spot using a cork borer size 4, and were mounted, live, in wells containing dH2O made in Carolina Observation Gel to enable diffusion of gasses. The abaxial of the leaf tissue was imaged using a Leica SP8 with 40x water immersion objective. Laser excitations for fluorescent proteins were used as described previously (Duggan *et al*., 2021).

### Membrane enrichment assays

Membrane enrichment was carried out by slightly modifying a previously described protocol (Abas & Luschnig, 2010). In brief, leaf material was ground to fine powder using liquid nitrogen and 2x volume of extraction buffer was added. Extraction buffer consisted of 0.81M sucrose, 5% (v/v) glycerol, 10 mM EDTA, 10 mM EGTA, 5mM KCl, and 100 mM Tris-HCl (pH 7.5) supplemented with 5 mM DTT, 1% Sigma Plant Protease Inhibitor Cocktail, 1 mM PMSF and 0.5% PVPP. After addition of the buffer, the samples were vortexed for a minute and the cell debris was cleared out by two subsequent centrifuges at 1000 xg for 5 min. The supernatant was diluted 1:1 using distilled water and an aliquot of the supernatant was separated as the total fraction (T). The remaining supernatant (200-300 μL) was further centrifuged at 21,000 xg for 90 min at 4°C. This centrifugation yielded the supernatant (soluble fraction, S) and membrane enriched pellet (membrane fraction, M). After separating the soluble fraction, the pellet was resuspended in diluted extraction buffer (without PVPP). All the fractions were diluted with SDS loading dye, and proteins were denatured by incubating at 50°C for 15 min. Western blotting was performed as previously described following SDS-PAGE. Endogenous plasma membrane ATPase was detected using anti-H+ATPase (AS07 260) antibody (Agrisera) as a marker to show the success of membrane enrichment.

## Supplementary information

**Figure S1:**
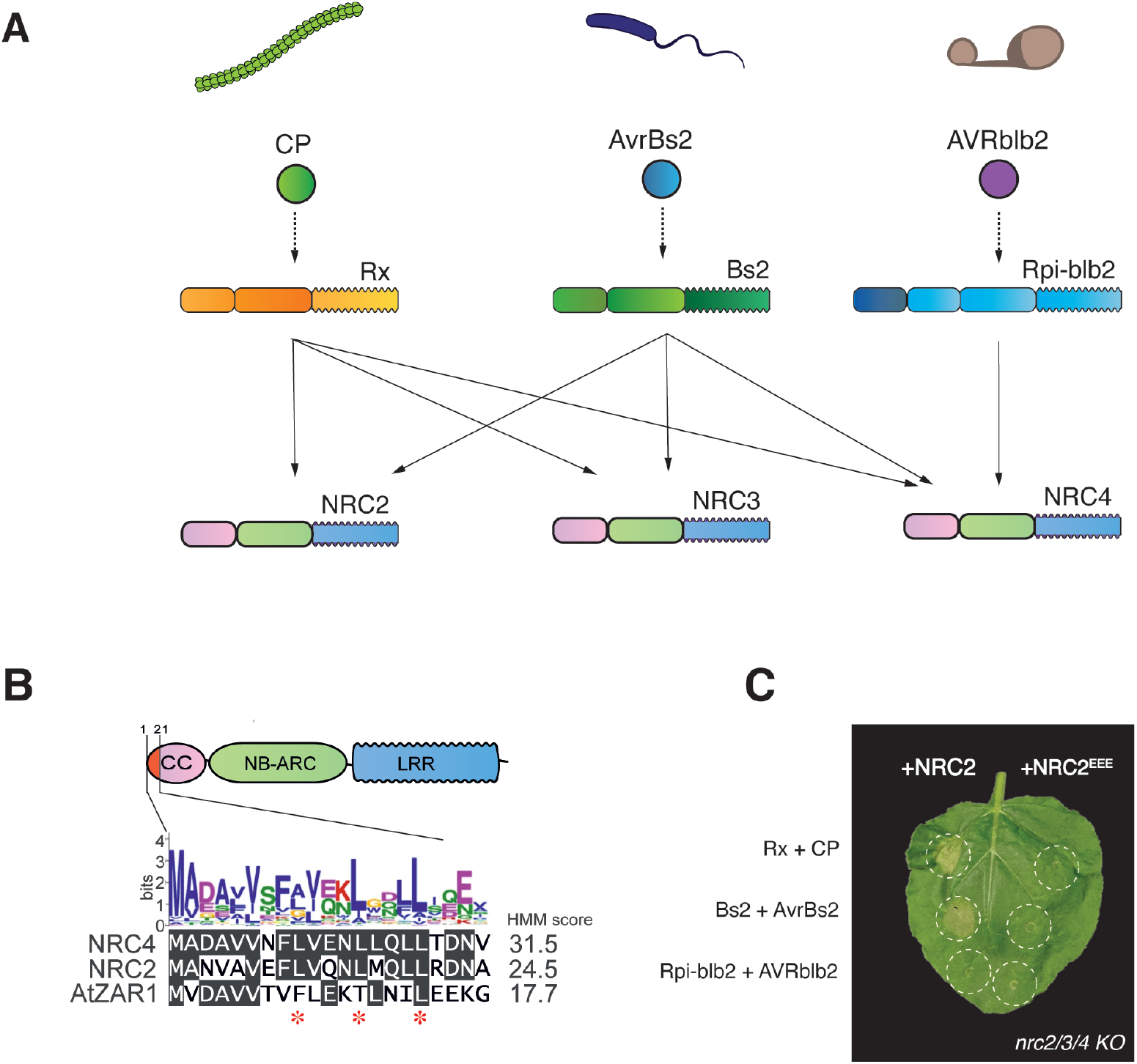
MADA motif mutants of NRC2 are unable to trigger cell death. (**A**) The Solanaceous NRC immune receptor network. Sensor NLRs such as Rx, Bs2 and Rpi-blb2 have specialized in detecting effectors from pathogens as diverse as viruses, bacteria and oomycetes. Sensors in the NRC network signal, sometimes redundantly, through their downstream helper NLRs, the NRCs. Different sensors exhibit different helper specificities. For example, Rx and Bs2 which recognize CP and AvrBs2 respectively, can signal through NRC2, NRC3, and NRC4. Rpi-blb2, which recognizes AVRblb2, can only signal through NRC4. Some sensor NLRs exhibit N-terminal extensions, represented in dark blue. (**B**) Much like AtZAR1 and NRC4, NRC2 has an N-terminal MADA motif. Alignment of NRC2, NRC4 and AtZAR1 N-terminal MADA motifs along with the consensus sequence pattern for the motif and the HMM score for MADA prediction of each sequence. Residues mutated in NRC2^EEE^ mutant are highlighted with red asterisks (positions 9, 13 and 17 respectively). (**C**) Unlike NRC2, NRC2^EEE^ does not complement Rx/CP and Bs2/AvrBs2-triggered hypersensitive cell death in leaves of *nrc2/3/4 N. benthamiana* CRISPR mutant lines. Representative leaves infiltrated with the appropriate constructs were photographed 5-7 days after infiltration. NRC2 and NRC2^EEE^ constructs are C-terminally 4xMyc-tagged. All effectors used are C-terminally GFP-tagged. All sensors used are C-terminally 6xHA tagged. One representative leaf is shown.

**Figure S2:**
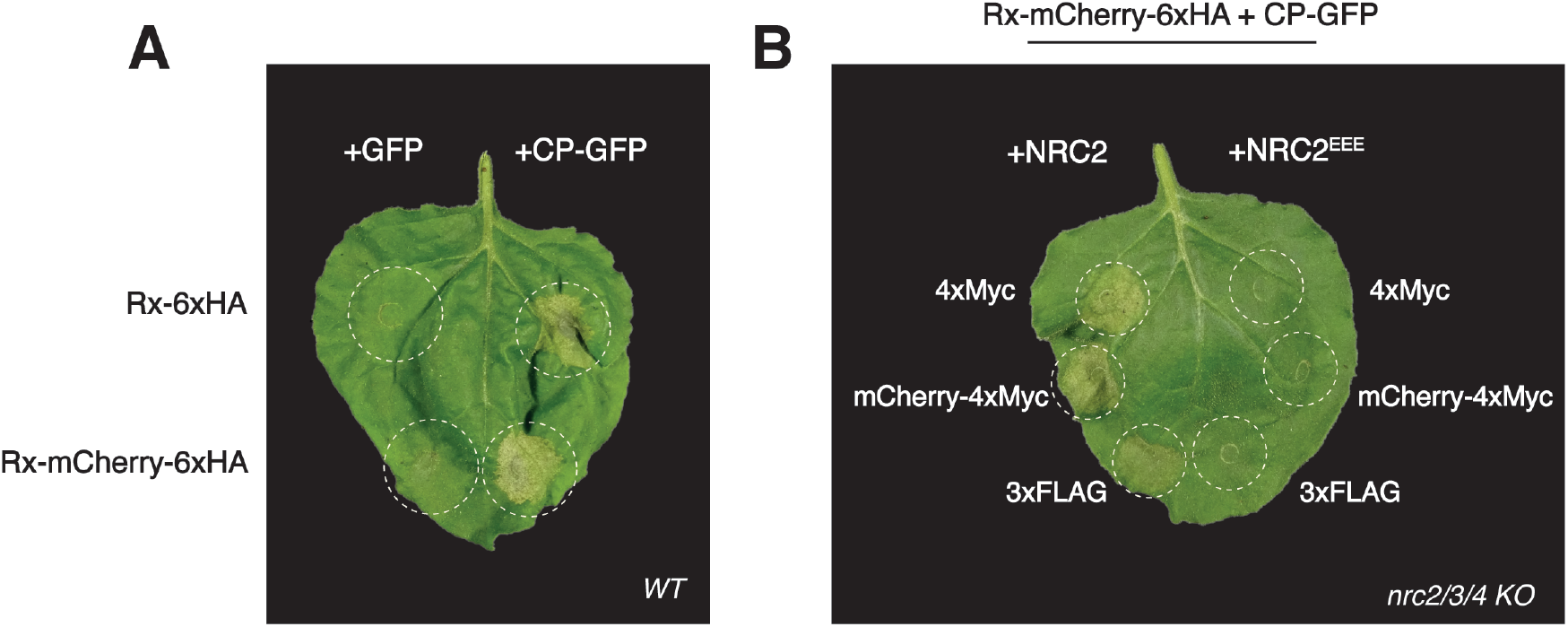
C-terminally tagged sensor and helper NLRs retain the capacity to trigger hypersensitive cell death. (**A**) Much like C-terminally 6xHA tagged Rx, C-terminally mCherry-6xHA tagged Rx can mediate hypersensitive cell death when activated by CP. Representative leaves of *WT N. benthamiana* were infiltrated with the appropriate constructs and photographed 5-7 days after infiltration. C-terminal tags are indicated. Free GFP (+GFP) was used as a negative control for C-terminally GFP-tagged CP (+CP-GFP). One representative leaf is shown. (**B**) Rx-mCherry-6xHA is compatible with all C-terminally tagged versions of NRC2 tested. Rx/CP-triggered hypersensitive cell death was complemented by C-terminally 4xMyc, mCherry-4xMyc and 3xFLAG variants of NRC2 respectively in leaves of *nrc2/3/4 N. benthamiana* CRISPR mutant lines when Rx was C-terminally tagged with mCherry-6xHA. The corresponding NRC2^EEE^ variants with the same C-terminal tag were no longer able to complement hypersensitive cell death. Representative leaves were infiltrated with the appropriate constructs and photographed 5-7 days after infiltration. One representative leaf is shown.

**Figure S3:**
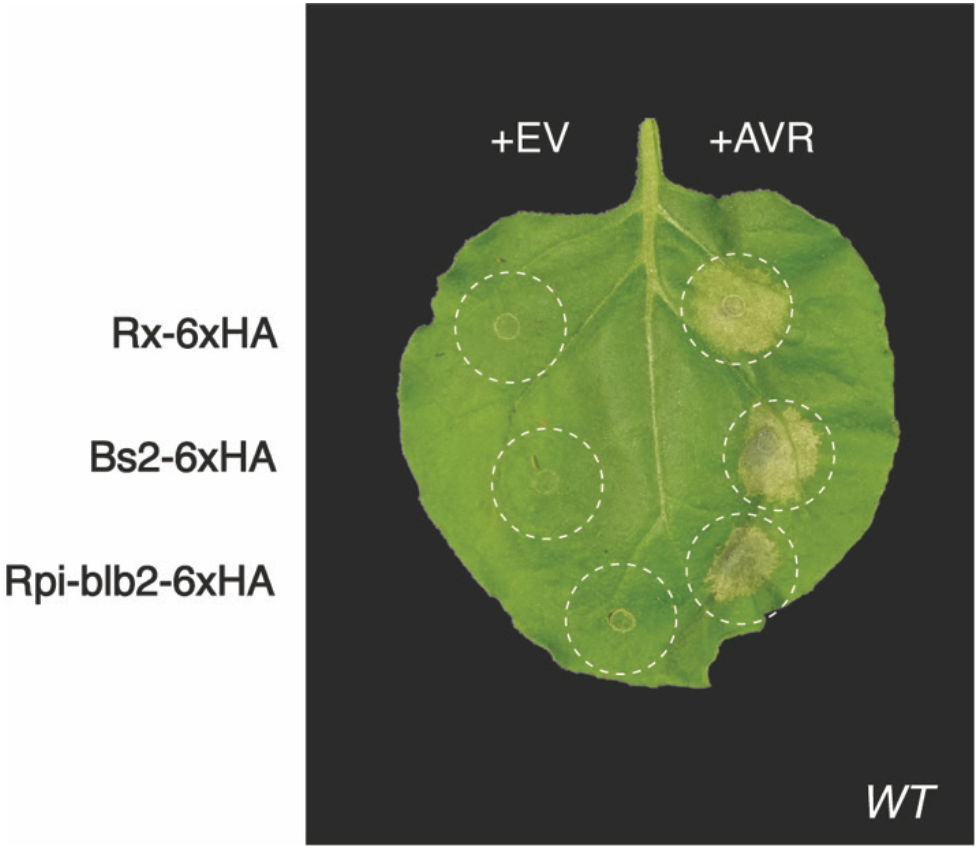
C-terminally 6xHA tagged sensor NLRs retain the capacity to mediate cell death. C-terminally 6xHA tagged Rx, Bs2 and Rpi-blb2 can mediate hypersensitive cell death when activated by CP, AvrBs2 and AVRblb2, respectively. Representative leaves of *WT N. benthamiana* were infiltrated with the appropriate constructs and photographed 5-7 days after infiltration. Free GFP was used as a negative control (EV) for C-terminally GFP-tagged effectors (AVR). One representative leaf is shown.

**Figure S4:**
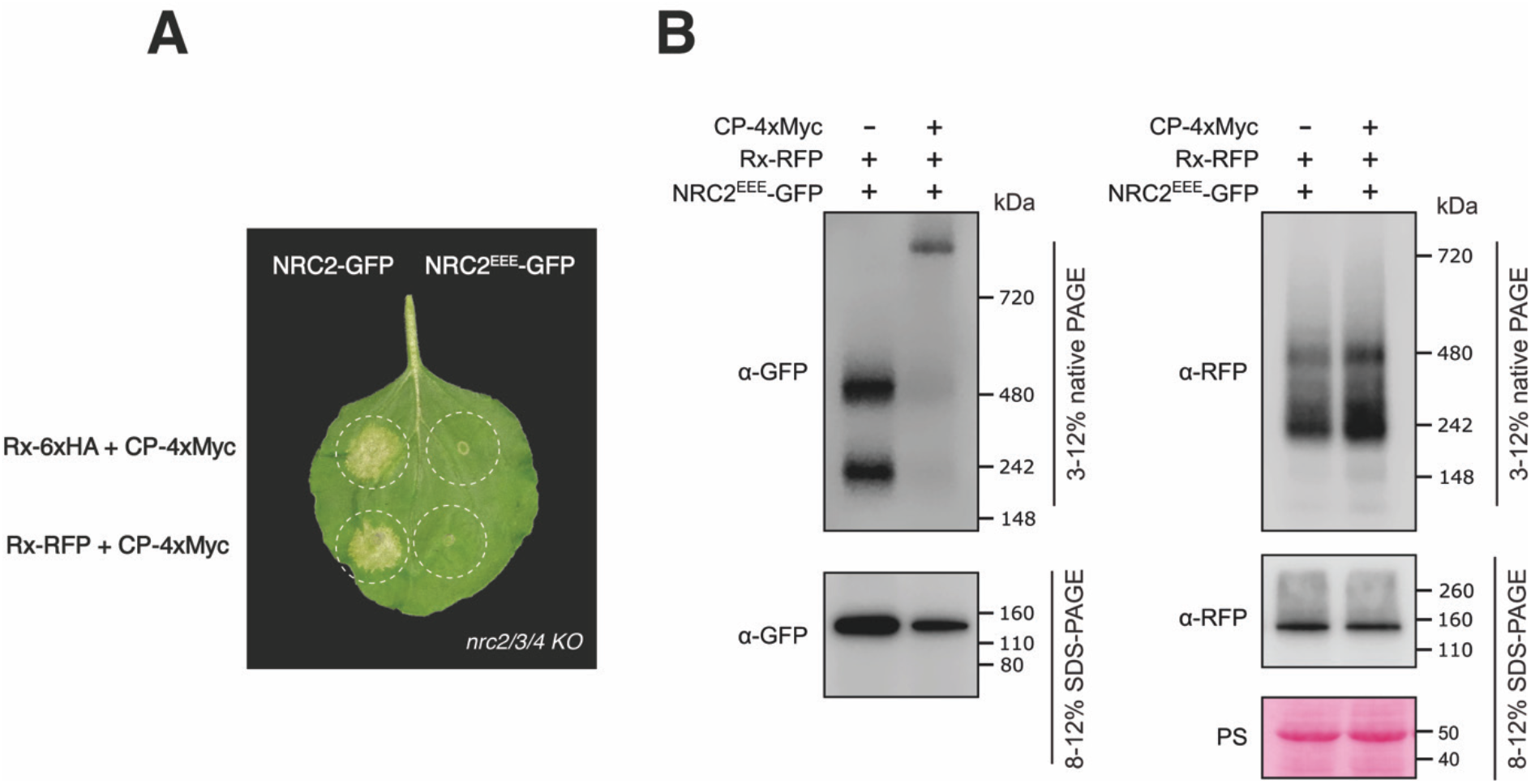
Fluorescent protein-tagged Rx and NRC2 retain cell death-mediating capacity and can correctly oligomerize upon activation. (**A**) C-terminally GFP-tagged NRC2 complements Rx/CP cell death in leaves of *nrc2/3/4 N. benthamiana* CRISPR mutant lines when Rx is C-terminally tagged with 6xHA or RFP. This cell death is not complemented with C-terminally GFP-tagged NRC2^EEE^. Representative leaves were infiltrated with the appropriate constructs and photographed 5-7 days after infiltration. One representative leaf is shown. (**B**) BN-PAGE and SDS-PAGE assays performed in parallel on protein extracts used for membrane enrichment assays with inactive and activated C-terminally RFP-tagged Rx and and C-terminally GFP-tagged NRC2^EEE^. Total protein extracts were run on native and denaturing PAGE assays in parallel and immunoblotted with the appropriate antisera labelled on the left. Approximate molecular weights (kDa) of the proteins are shown on the right. Rubisco loading control was carried out using Ponceau stain (PS). The experiment was repeated 2 times.

**Movie S1:**

**Movie S1: NRC2 forms PM-associated puncta upon activation by *Potato virus X* coat protein and Rx**.

3-D movies of inactive (left) or CP-activated (right) NRC2^EEE^-GFP (shown in yellow) co-expressed with Rx-RFP (not shown) in leaves of *nrc2/3/4 N. benthamiana* CRISPR mutant lines. CP was C-terminally 4xMyc-tagged. Free 4xMyc tag was used for the inactive negative control.

**Table S1:**

**Table S1: List of primers and constructs used in this study**.

## Acknowledgements

We are very thankful to several colleagues for discussions and ideas. We thank Daniel Lüdke (The Sainsbury Laboratory, Norwich, UK) for valuable comments on this paper. We thank all members of the TSL Support Services for their invaluable assistance. We thank S. Hrek. M.C. and S.K. dedicate this paper to the memory of Diego A. Maradona. A.V.C. was funded by the John Innes Foundation and TSL. The Kamoun Lab is funded primarily from the Gatsby Charitable Foundation, Biotechnology and Biological Sciences Research Council (BBSRC, UK), and the European Research Council (BLASTOFF).

## Author Contributions

M.P.C: Conceptualization, Methodology, Validation, Formal Analysis, Investigation, Data Curation, Writing – Original Draft, Writing – Review & Editing, Visualization, Supervision, Project Administration.

H.P.: Methodology, Validation, Formal Analysis, Investigation, Data Curation.

Y.T.: Methodology, Validation, Formal Analysis, Investigation, Data Curation, Writing – Review & Editing, Visualization.

C.D.: Methodology, Validation, Formal Analysis, Investigation, Data Curation, Writing – Review & Editing, Visualization.

E.L.H.Y: Validation, Investigation, Data Curation, Writing – Review & Editing, Visualization.

A.V.C.: Validation, Formal Analysis, Investigation, Data Curation, Writing – Review & Editing.

J.K.: Resources, Writing – Review & Editing.

H-K.A.: Conceptualization, Methodology, Writing – Review & Editing.

C-H.W.: Conceptualization, Methodology, Writing – Review & Editing.

T.O.B.: Resources, Supervision, Funding Acquisition. Project Administration.

L.D.: Conceptualization, Resources, Supervision, Writing – Review & Editing, Project Administration.

S.K.: Conceptualization, Resources, Supervision, Funding Acquisition, Visualization, Writing – Original Draft, Writing – Review & Editing, Project Administration.

